# Evolution of vocal production learning in parrots

**DOI:** 10.1101/2024.11.05.622162

**Authors:** A Krasheninnikova, SQ Smeele, M Snijders, E Haldar, J Carpenter, R Zamora, M Naguib, JBW Wolf, M Gahr, AMP von Bayern

## Abstract

Vocal production learning (VPL), the capacity to imitate sounds, is a crucial, but not exclusive component of human language. VPL is rare in animals but common in birds, where it evolved independently in songbirds, hummingbirds, and parrots. Parrots (Psittaciformes) learn new vocalizations throughout their lives and exhibit astonishing vocal flexibility and imitation capacity. They can copy allospecific sounds, e.g., human words and learn their associated meanings. Parrots, therefore, present an intriguing model to shed light on how VPL evolved and how it may relate to other language-relevant traits. How widely VPL is distributed across Psittaciformes and to what extent (qualitative) species differences exist, remains unknown. Here, we provide the first comprehensive overview of the phylogenetic distribution of (allospecific) VPL in this clade by conducting surveys of publicly available video footage. Out of the 398 currently recognized extant species, we found videos for 163, of which 136 showed evidence of VPL. Phylogenetic analyses suggest secondary losses and reacquisitions of VPL covarying with socioecological parameters (gregariousness), life-history (longevity), and morphological (body size) traits. This study provides the first insights into interspecific variation in vocal learning across all parrot species and reveals potential socio-ecological drivers of its evolution.

**Significance:** Little is known about the selective forces that favor the evolution of vocal production learning (VPL), a rare trait in animals and a prerequisite for the evolution of human language. We provide the first insights into interspecific variation in VPL in the evolutionary history of parrots and uncover candidate evolutionary drivers. The current data suggest that the evolution of VPL within parrots has been highly dynamic, with multiple secondary losses and reacquisitions. Our model showed that VPL most likely was the ancestral state. Sociality, longevity and body size explain variation in VPL together with a highly uncertain effect of brain size. The findings may motivate comparative work in other taxa and contribute to research into the evolutionary origins of human language.

## Introduction

Vocal production learning (also referred to as imitative learning or vocal mimicry) is the capacity to learn vocalizations outside of the ”innate” vocal repertoire(1), and is a prerequisite for human spoken language (2). Without the ability to generate and acquire a large, open-ended vocabulary, the flexible, expressive power of language would be very limited (3). Vocal production learning (VPL hereafter) is a rare trait found in only five distantly related groups of mammals (humans, bats, cetaceans, pinnipeds, and elephants) and three distantly related groups of birds (songbirds, hummingbirds, and parrots). Given the scarce evidence for VPL in non-human primates (4), research has focused on songbirds as the dominant model for studying the developmental processes, mechanisms of acquisition, and evolutionary history of VPL over the past few decades(3–5). While Petkov and Jarvis (4) suggest that striking neurogenetic underpinnings of VPL have convergently evolved in humans and songbirds, this interpretation is contentious. Critics argue that the parallels may be superficial and call for more detailed comparative studies to substantiate claims of convergent evolution (e.g., (6). Despite these debates, parallels between song development of birds and human language development have been suggested, as reviewed by Aamodt et al. (7). For example, most vocally learning songbirds acquire their ability to sing similarly to how infants learn to speak, i.e., going through sensitive learning and “babbling” (i.e., practicing) phases (5, 8, 9).

Parrots are one of the most versatile avian vocal production learners (10–12) and, in fact, resemble humans more in their VPL traits than songbirds (1, 13, 14). Like humans, most parrots are open- ended vocal learners, i.e., their VPL is not limited to a particular sensitive period like in many songbirds, but they modify their vocal repertoires throughout life (13). Moreover, in parrots, both sexes use learnt vocalizations, rather than just males, as is the case in most songbirds (13). Another crucial aspect is that parrots use their learnt vocalizations in a wide variety of social contexts (7, 13, 14) rather than primarily for territorial defense or mate attraction (1), even though song also functions beyond those contexts (15). Recent research revealed that parrots also possess developmental babbling phases analogous to human prelinguistic development (16). In addition, parrots share several general life history traits with humans, such as longevity, slow development or extended parental care, and also live in complex, individualized societies (13). Finally, they stand out among vertebrates for their large relative brain sizes (17, 18), high neuron densities (19) and remarkable socio-cognitive abilities that make them more prone to complex communication (see (20, 21) for reviews). These similarities between parrots and humans make them a suitable model for studying VPL evolution, possible origins, and selection pressures for language-relevant traits (13).

Parrots are also well known for their ability to imitate human speech and represent one of the few animal taxa in which referential signaling has been demonstrated (22). Alex, the famous African grey parrot *(Psittacus erithacus*), learned over a hundred human words which he associated with meaning (23). A possible reason why parrots can mimic human speech is that their acoustic range of auditory perception and call production appears to overlap with that of humans (12, 24–27). Like humans, they also use tongue and beak movements for vocal articulation (8, 12, 28). The capacity of parrots to imitate sounds of human speech well outside the species-typical repertoires provides an opportunity to study VPL capacity (29) and inter-specific variation therein (1).

The most likely function of VPL in parrots is to enable social communication within flocks, for example, when mediating flock exchanges during social foraging (13, 30). Parrots have one or multiple types of contact calls designed explicitly for social communication and, in some species, contain clear individual signatures (e.g., (30, 31). There is even evidence of a voice print going across call types, enabling individuals to remain recognizable, even during flexible exchanges (32). Contact calls can also be modified in social groups. For example, budgerigars (*Melopsittacus undulatus*) converge on the call of conspecifics (33) leading to additional social signatures, and spectacled parrotlets (*Forpus conspicillatus*) may use distinct contact calls (34) to selectively ‘address’ specific individuals directly. It has been pointed out that the complex social systems of many parrot species, their highly complex and dynamic social fission-fusion foraging cultures (considered a result of their extraordinarily challenging diet of toxic, unripe seeds), and their possible dependency on social learning may select for acoustic individual recognition, vocal communication and exchange of information (1, 13). Consequently, many aspects of the socioecology and life history of different parrot species may account for possible inter-specific variation in VPL capacity across parrots. For example, continuous VPL may be vital for species with high longevity (35), fluid social dynamics and social foraging (13, 36). Furthermore, parrots vary enormously in body mass, a morphological trait that can shape some acoustic features of parrots’ vocalizations (e.g., peak frequency and duration (37)) thus leading to various levels of vocal plasticity in differently sized species. Finally, the recently discovered shell regions unique to the parrots’ brain tend to be larger in larger-brained species that are also typically considered to have more advanced vocal and cognitive abilities (10). Therefore, parrots as open-ended vocal learners and highly diverse avian taxon with 398 extant species that vary in body size (between 10g, *Microspitta pusio*, and 2250g, *Strigops habroptilus*, (38), life history parameters and socio-ecological variables, represent an ideal model system for studying the evolution of VPL. Given that the most striking evidence for VPL in parrots comes from individuals kept as pets mimicking human speech or other anthropogenic sounds from their environment (39), and considering that the majority of parrots are kept as pets or in zoos, a focus on parrots kept in human care may provide a fruitful first venue for research on vocal mimicking capacities across parrot species.

In this study, we took advantage of the social media platform YouTube to gain first insights into the phylogenetic distribution of and interspecific variability in VPL of parrots. We examined the ability of all 398 parrot species to imitate human speech and anthropogenic sounds and thus the yet untested prediction that vocal learning is ubiquitous among all parrots and evolved at the base of the Psittaciformes radiation 29 Mio years ago (10, 40).

In order to examine potential evolutionary drivers, we considered four variables that we predicted to select for increased VPL (longevity, social complexity, body size, and relative brain size) and made the following specific predictions for each of them. Parrots learn new vocalization throughout their life, therefore, long life spans may entail more opportunities for vocal learning (1). Also, longer-lived species are more likely to face social and environmental changes, which may select for greater flexibility in communication. We thus expected longer-lived species to be more likely to exhibit VPL or to be better at VPL than shorter-lived species. Furthermore, we predicted that they would exhibit larger learnt vocal repertoire sizes than shorter-lived species. Similarly, complex social dynamics may select for VPL, e.g., by facilitating mediation during flock exchanges (1). We took gregariousness (defined as nesting close together or being colonial) as an available proxy for social complexity and expected highly gregarious species to exhibit (better) VPL more likely but also to show larger learnt vocal repertoire than less gregarious species. Furthermore, larger parrots produce lower fundamental frequencies with longer wavelengths than smaller parrots; thus, we expected larger species to show a higher range of imitations of human words. Finally, vocal learning is often considered a cognitively demanding capacity that requires a relatively high brain processing capacity ((3, 41), but see (42)). Moreover, relative brain size is correlated with key features that support VPL, such as increased longevity, which provides a longer time span for learning and utilizing complex vocalizations (43). Additionally, it is linked to the unique neuroanatomical features of the parrot brain, particularly the shell structures. Research by Chakraborty and colleagues (10) revealed significant plasticity in the relative sizes of song nuclei among parrot species, with the shell regions being larger in species with larger brains, which are typically considered to possess more advanced vocal and cognitive abilities. This pattern demonstrates that as brain size increases, the relative size of shell regions also increases, albeit slower, suggesting a complex relationship between brain size and vocal learning capacities. Thus, we expected larger-brained species (relative to body size) to be better at VPL.

## Results

We collected data on the occurrence of VPL in all 398 extant parrots and found videos for 163 species, of which 162 could be considered to be in human care (i.e., they were either reported in international trade by CITES and/or found in captivity, see Tab. 1 and Supplementary Information A, for more explanation and an overview). In total, 136 species showed evidence of VPL, i.e., produced at least one mimicked allospecific vocalization. For the remaining 235 species no videos were found, but 192 of these species were kept in human care. Assuming that the capacity to mimic human speech or anthropogenic sounds is sufficiently conspicuous to be captured on video when exhibited by a human companion, the absence of evidence of VPL in the 192 species kept in human care was, with all due caution, scored as tentative evidence of absence for VPL (see Methods and Discussion). In contrast, the 43 species not kept in human care were treated as missing data points (regarding VPL capacity of these species).

**Table 1:** Overview of the number of species for which YT videos were present or absent, species that were reported in human care, and for which VPL ability was present/absent, or data was lacking.

| YT videos<br>163 species |  |  |  | No YT videos<br>235 species |  |  |
| --- | --- | --- | --- | --- | --- | --- |
| In human care<br>162 |  | NOT in human care<br>1<sup>*a</sup> |  | In human care<br>192 |  | NOT in human care<br>43<sup>a</sup> |
| VPL<br>135 | No VPL<br>27 | VPL<br>1 | No VPL<br>0 | VPL<br>0<sup>b</sup> | No VPL<br>192<sup>b</sup> | Missing data<sup>a</sup> |

<sup>a</sup>These species are excluded from the evaluation of the effects of longevity, relative brain size, sociality and body size on the likelihood of a species exhibiting VPL.
<sup>b</sup>Since the absence of evidence in the species that occur in human care (i.e., in international trade and/or captivity) appears reliable, as the ability to mimic anthropogenic sounds or human speech is likely to be captured on video if it occurs, we did not exclude these species when assessing the effects of longevity, relative brain size, sociality and body size on the likelihood of a species exhibiting VPL. However, we also re-ran the analyses with only the subset of species for which we found YT videos. The effects found remained stable for this subset of data as well.
<sup>\*</sup>*Prioniturus verticalis*

The ancestral state reconstruction based on the available data revealed an early presence of VPL in parrots (for the full distribution, see Figure 1 and Supplementary Information B, Fig.SB1). There was a strong phylogenetic signal, with closely related species showing a similar expression of VPL. The signal was much weaker further down the tree with species that diverged about 10 mya or more having very low covariance (see Supplementary Information B, Figure SB2). We ran four phylogenetically controlled models to test the total effect of longevity, relative brain size, sociality, and body size on the probability that a species possessed VPL (excluding the 44 species which were not reported in human care). All variables had a positive total effect, although the effect of relative brain size was highly uncertain (longevity: mean = 0.83, 89% PI = (0.40, 1.28); relative brain size: mean = 0.64, 89% PI = (-0.44, 1.68); sociality: mean = 0.78, 89% PI = (0.05, 1.50); body size: mean = 0.93, 89% PI = (0.65, 1.24); also see Figure 2).

**Figure 1:**
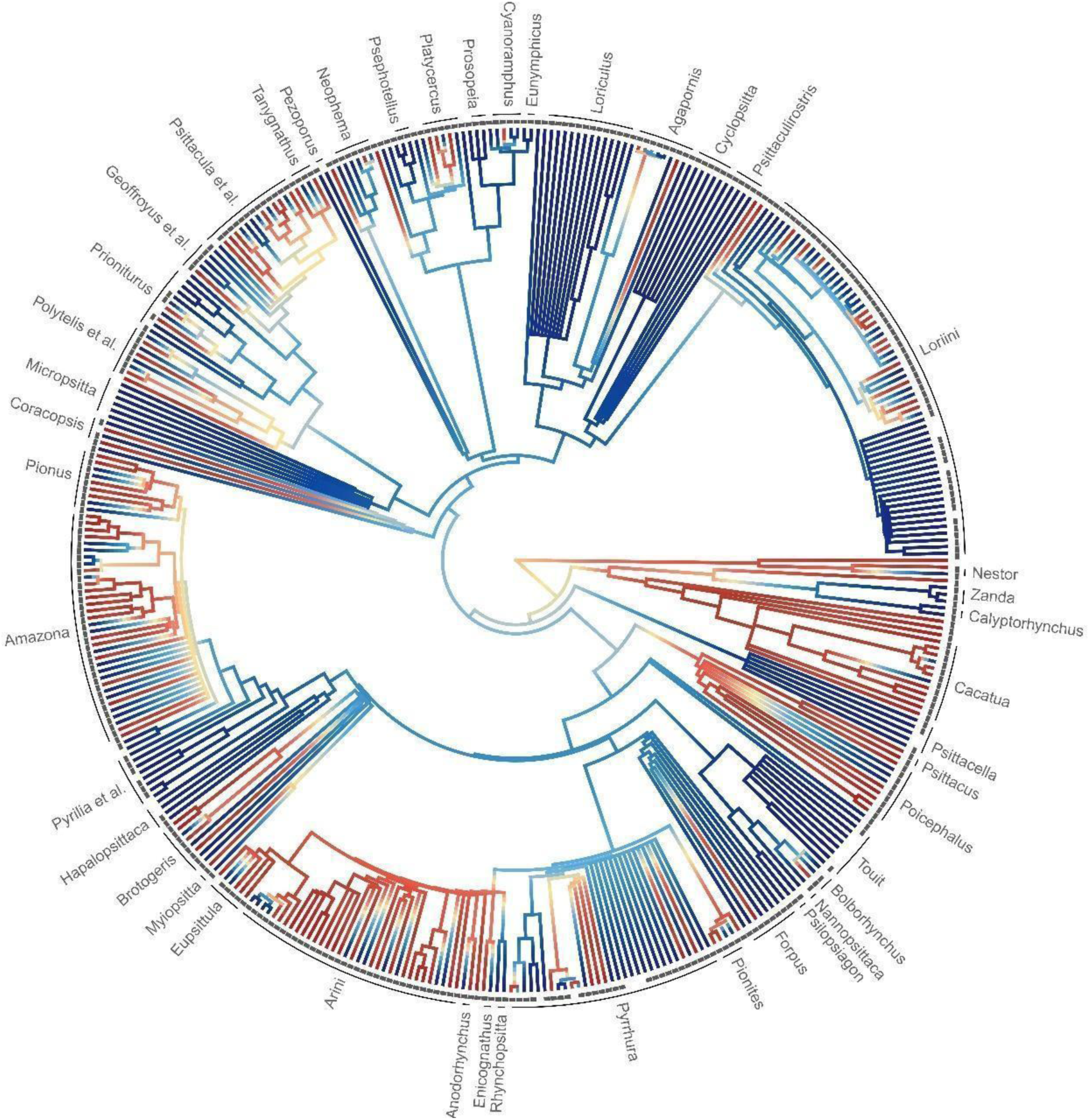
Phylogenetic tree of all parrot species indicating the presence and absence of VPL ability and species for which VPL data are lacking. Colors represent the reconstructed state of VPL of ancestral lineages ranging from blue = no VPL detected to red = VPL detected. The grey squares indicate the species reported to be in human care. Missing grey squares indicate the 44 species not reported in human care. Note that for 43 out of these 44 species, VPL ability remains unknown (=missing data points).

**Figure 2:**
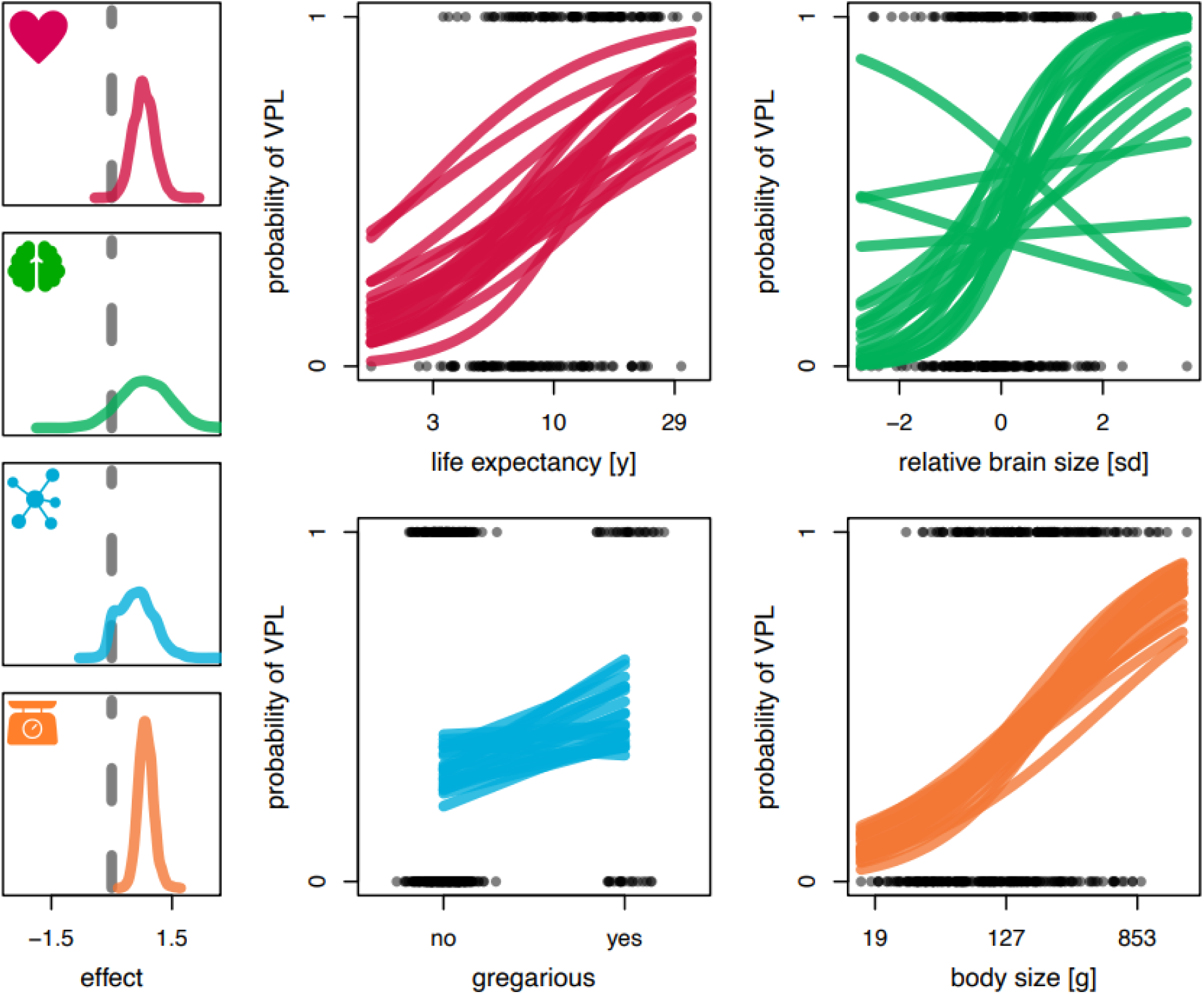
Variables influencing a species’ probability of exhibiting VPL (excluding 44 species not reported in human care). Lefthand side: posterior densities of the effect of longevity (red), relative brain size (green), gregariousness (blue), and body size (orange). For gregariousness, the contrast between a non-gregarious and a gregarious species is shown. For all other variables, the slope is shown. Righthand side: scatterplots of the raw data (grey) and 20 posterior predictions (colored lines) per variable.

Next, we recorded “VPL total repertoire size” (i.e., scored as the number of distinct mimicked vocalizations an individual produced, see methods and Supplementary Information A for details) for 843 individuals across 136 species (i.e., between a minimum of 1 and up to 25 individuals per species). Again, we ran four phylogenetically controlled models to test for the total effect of longevity, relative brain size, sociality and body size on repertoire size. Longevity had no effect, relative brain size had a highly uncertain effect, while sociality and body size had small positive effects (see Figure 3). The within-species variability of the VPL total repertoire size was low (Supplementary Information B, Figure SB3).

**Figure 3:**
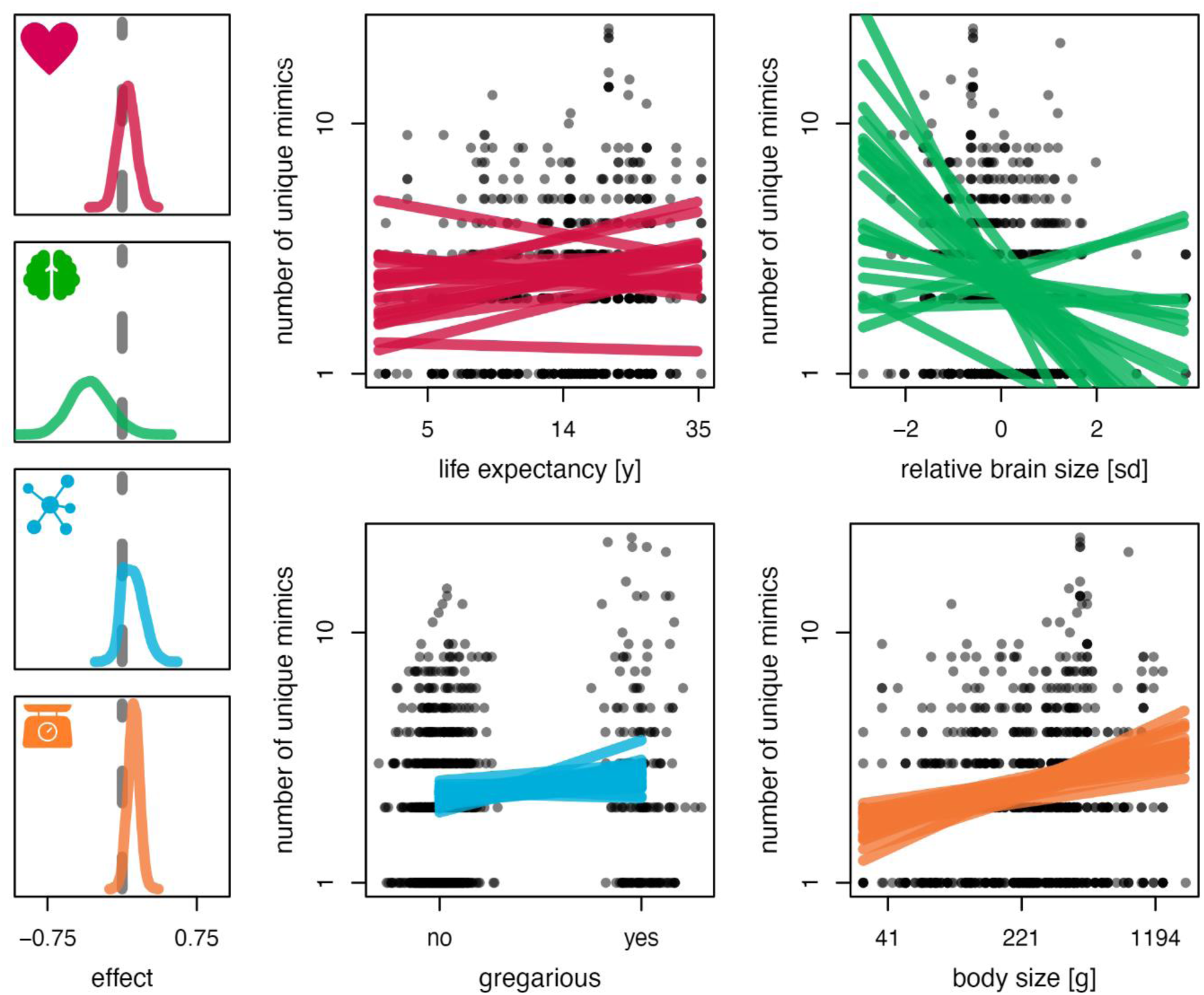
Variables influencing the number of distinct mimicked vocalizations an individual produced (i.e., VPL total repertoire size). Lefthand side: posterior densities of the effect of longevity (red), relative brain size (green), gregariousness (blue) and body size (orange). For gregariousness, the contrast between a non-gregarious and a gregarious species is shown. For all other variables, the slope is shown. Righthand side: scatterplots of the raw data (grey) and 20 posterior predictions (colored lines) per variable.

Last, we tested the influence of the four variables on a measure of “VPL anthropophonic repertoire size” (i.e., measured as the number of distinct recognizable human words an individual produced in a video; see Figure 4). Longevity had a clear total effect. Relative brain size had no clear effect. Sociality and body size had small and uncertain effects. In contrast, an analysis of qualitative measures of the “VPL quality” (i.e., scored as how recognizable the mimicked vocalizations were: low - babbling, no clear words or melodies recognizable, medium - words/melodies recognizable but not highly defined, high - mimicked words/melodies very well defined and recognizable) revealed a small effect of longevity and strong effect of body size (Supplementary Information B, Table SB1).

**Figure 4:**
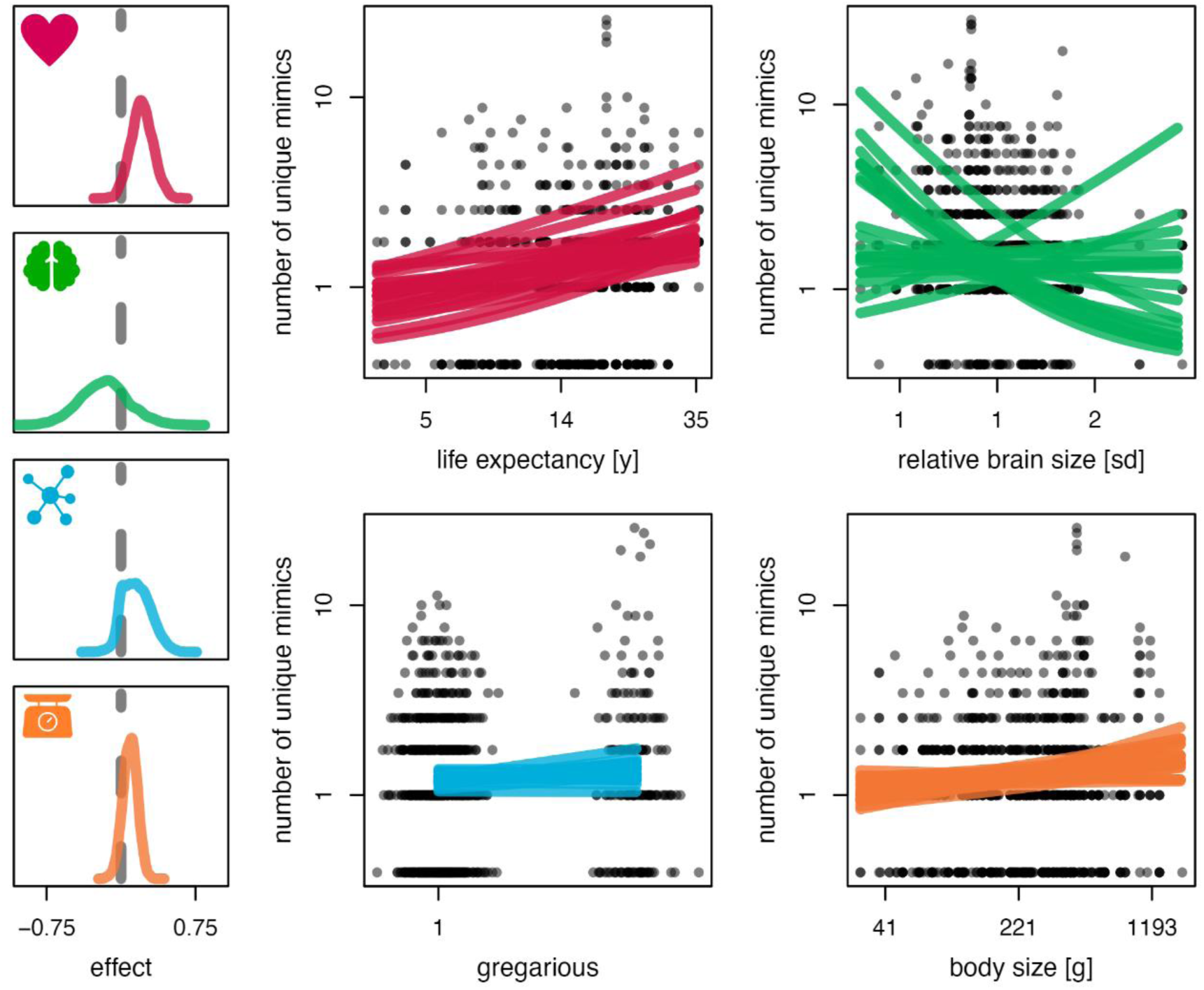
Variables influencing the number of distinct human words an individual produced (i.e., “VPL anthropophonic repertoire size”). Lefthand side: posterior densities of the effect of longevity (red), relative brain size (green), gregariousness (blue) and body size (orange). For gregariousness, the contrast between a non-gregarious and a gregarious species is shown. For all other variables, the slope is shown. Righthand side: scatterplots of the raw data (grey) and 20 posterior predictions (colored lines) per variable.

## Discussion

### The phylogenetic distribution of vocal learning in parrots

This study reveals an early presence of VPL among *Psittaciformes*. We show that the kaka (*Nestor meridionales*), one of the most basal parrot species, is capable of VPL. The finding that VPL capacity is present in this very basal group of parrots suggests that it probably derived from a common ancestor of all parrots. This is in line with previous studies on the neural underpinnings of VPL in parrots, suggesting that the specialized brain areas associated with vocal learning and song production (song nuclei) have evolved early on in parrot diversification, at least 29 million years ago, before the genus *Nestor* split from the other parrot lineages (10). The phylogenetic map of VPL we obtained not only suggests that VPL is ancestral in parrots. It also uncovers intriguing evolutionary scenarios for vocal production learning (VPL) within the Psittacoidea superfamily, offering two potential narratives. Firstly, it is possible that the superfamily *Psittacoidea* lost VPL secondarily, meaning that the great majority of VPL species (i.e., 122 species in *Psittacoidea*) actually exhibit a derived, loss-of-function trait (44).

Alternatively, the pattern might indicate that VPL is not originally ancestral within Psittacoidea, with the trait having arisen independently multiple times across various species. The frequent independent emergence of VPL under this scenario would suggest a strong adaptive advantage in certain ecological and social environments, counterbalanced by many secondary losses where the evolutionary costs of maintaining VPL outweigh its benefits (36)(Sewall 2015). Note that even though secondary losses of social behavior are quite rare in animals, they have been noted in birds previously (e.g., reversed solitary roosting in communally roosting clades, (45)Beauchamp 1999). In this context, the functional learning of vocalizations and species recognition are crucial aspects that merit further exploration. When the function of VPL is no longer adaptive, evolutionary pressures might favor the suppression of this complex capability, allowing individuals to revert to simpler, more recognizable calls.

This point underscores the complex interplay between genetic, ecological, and social factors that shape the evolution of vocal learning in parrots. It suggests that the adaptive value of VPL is intricately linked to the specific social and environmental demands faced by a species. As such, understanding the functional implications of VPL in different ecological contexts is essential for elucidating the reasons behind its evolutionary persistence or loss in certain lineages.

Hence, according to the first scenario, an unexpected evolutionary pattern arises; the two superfamilies *Strigopoidea* (3 sp.) and *Cacatuoidea* (21 sp.) apparently kept the ancestral VPL, whereas the much bigger superfamily *Psittacoidea* (True parrots, 350 sp.) lost VPL, except for the sub-family *Psittacinae* of afrotropical parrots (12 sp.; sister group to the neotropical *Arinae*). In some *Psittacoidea* genera (e.g., in *Forpus, Pyrrhura)*, the majority of species lack evidence for VPL, whereas a few species exhibit VPL (in *Forpus*, 1 out of 8 species, and in *Pyrrhura*, 10 out of 31 species). This indicates that those few species seem to have newly and possibly independently evolved VPL (reversal) which their ancestors had lost according to the current data. However, it is important to consider the distinction between the independent evolution of a trait and the reactivation of a previously existing trait. Analogous to flight in certain flightless birds, which might more readily re-evolve due to the pre-existence of wings, parrots likely retain much of the necessary brain structure for VPL even after millions of years without active use of the trait. This residual capacity suggests that reactivating VPL in parrots could be less challenging than evolving such complex behaviors de novo in other species lacking any foundational brain structures for vocal learning. In other Psittacoidea groups, e.g., in all genera from the neotropical Arini tribe, (except for the *Pyrrhua* genus), most species exhibit VPL, while few species lack it. For example, all eight species belonging to the *Ara* genus exhibit VPL, whereas 2 out of 6 Eupsittula species, 2 out of 6 *Aratinga* species lack VPL. This suggests in turn that in this taxonomic group VPL has re-evolved (reversal) but then was lost a third time in several species. Tertiary loss of a trait in evolution is only known from eye evolution in cave-dwelling organisms (44).

On the other hand, in the Loriini group, the Lorius and Glossopsitta branches appear to have VPL but that some have secondarily lost it, advocating for a second scenario that assumes losing VPL is more likely than (re-)evolving VPL. Our main result suggests the absence of VPL in the last common ancestor of Lorinii, but we found also that their two closest relatives have VPL. It could therefore also be that VPL was present in the last common ancestor and was lost multiple times in Lorinii, due to the high cost it incurs.

Potential costs of VPL include developmental costs of the brain tissue due to the need for increased cognitive capacity (10, 41) and opportunity costs stemming from the need for social feedback essential for learning (36). Many open-ended vocal learners such as elephants, cetaceans, and humans exhibit extended juvenile and adolescent phases along with prolonged parental care (1). This extended period facilitates crucial learning from parents and other conspecifics (46, 47), as seen in human infants whose prelinguistic development is significantly shaped by caregiver interactions (48). However, these prolonged care periods can impose substantial costs on parents by delaying the time their offspring become communicatively independent and capable of reproducing (49). Another potential cost is that hybridization risk can increase if vocalizations become less stereotyped, since species cannot reliably use species specific vocalizations to recognize conspecifics for mating (50, 51). Finally, while vocal learning allows for the rapid spread of new vocalizations, meaning might be lost if vocalizations spread too quickly. Referential calls, such as the alarm calls of vervet monkeys are likely innate, since alarm calls need to be recognized immediately, and there is no time for social learning (52). The developmental constraints may limit the evolution of VPL to species with longer lifespans (1), and then only to species that gain a clear advantage from VPL. Yet, given the wide occurrence of VPL across parrots, including the many species that have secondarily lost and re-evolved VPL according to our current data, it seems to be highly adaptive in parrots.

Our study strategically utilized proxies for vocal production learning (VPL), which are recognized as reliable despite inherent challenges. A notable concern might be the misclassification risk, particularly the assumption that the absence of VPL in YouTube videos equates to a genuine absence of VPL capabilities. To counteract this, we enhanced our methodological approach by also analyzing a dataset exclusively comprising individuals in human care (Supplementary Information B, Figure SB4). This subset is likely more consistently represented on YouTube, thereby reducing bias related to VPL visibility. The consistency between findings from both datasets underscores the robustness and reliability of our results. Additionally, we acknowledged the challenges in accurately assessing sociality and longevity due to the limited documented socio-ecological backgrounds of many parrot species (53). In response, we utilized the best available proxies (e.g., longevity data, (43), while remaining cautious of their approximative nature. This careful consideration ensures a more informed interpretation of how these factors might influence the evolution and expression of VPL, recognizing the varied social structures and lifespan across parrot species which correlate with cognitive capacities and complex behaviors like VPL. Finally, our use of publicly available online videos, while presenting its unique challenges, offers substantial benefits over other citizen science methods such as questionnaires, which can suffer from phrasing biases (54). By employing crowdsourcing on social media platforms, we were able to apply objective criteria and directly measure behaviors like frequencies and durations (55).

Overall, our findings suggest that VPL has evolved and re-evolved within *Psittaciformes* multiple times making them an even more interesting model system. More comprehensive data, including a better knowledge of the different species’ socio-ecology and further detailed analyses will eventually reveal the underlying evolutionary dynamics. Some groups, e.g*., Amazona* (18 VPL vs 16 non-VPL species), *Pionus* (5 VPL vs 4 non-VPL species), and *Psittacula* (7 VPL vs 5 non-VPL species) genera exhibit substantial variation and will be key to understand the evolutionary forces that select for or against VPL in better resolution. It needs to be considered that VPL is a complex behavioral phenotype type likely involving multiple components (56). More detailed studies of the possibly very different VPL phenotypes in different taxonomic groups of Psittaciformes, combining genetic, neurological, developmental, and behavioral approaches with socio-ecology, will shed further light on VPL evolution.

### Possible evolutionary drivers of VPL ability and VPL repertoire sizes in parrots

Examining potential evolutionary drivers selecting for VPL in parrots, we found that longevity, sociality, and body size best predicted the species differences in VPL ability: as predicted, the probability that a species possesses VPL ability was generally higher for long-living, large and gregarious species. As outlined earlier, long-lived species often have a prolonged developmental period that is crucial not just for exposure to different vocal models but also for the extensive practice needed to master VPL (1, 57). Our finding that longer-lived species were more likely to possess VPL and showed higher “VPL anthropophonic repertoire size” compared to shorter-lived species is in line with the assumption that VPL may be constrained to long-lived species.

Interestingly, no effect of longevity on VPL total repertoire size was found. This might indicate that although parrots are open-ended vocal learners, they may not just expand their repertoires over time like some songbirds do (58), but rather, and similarly to hummingbirds (59), “replace” some parts of it with new vocalizations/adjust to constantly changing socially transmitted group-specific vocalizations. This does not necessarily imply that vocalizations are “overwritten” but that some sounds are no longer relevant and need to be updated when socioecological conditions change or when group-specific vocal traditions are transformed via cultural processes. Given the potential function of learned vocalizations in mediating social interaction and acceptance into newly formed groups in ever changing social structures with fission-fusion dynamics, it appears reasonable that sociality but not longevity affects the repertoire size of learned vocalizations (60). These findings align with the results recently reported by Benedict and colleagues (29). The authors found that companion grey parrots expanded their vocal repertoires through the juvenile phase but did not show reliable expansion among adults, suggesting that the vocal repertoires of adult companion parrots may change by element replacement rather than addition (29).

In contrast, there was a clear effect of longevity when looking only at the number of distinguishable human words learnt (“VPL anthropophonic repertoire size”). The correlation between body size and longevity has been recently confirmed for six taxonomic classes, including birds (Kuparinen et al. 2023), and was accounted for in our statistical analysis (see Supplementary Information C, **Figure SC1** Directed Acyclic Graph). This aligns with previous findings by Benedict et al. (29), showing that - according to a questionnaire survey conducted on parrot owners - companion parrots’ repertoire size was the largest for human words. In their natural habitat, parrots imitate socially significant vocalizations such as the individually distinct contact calls of certain group members (e.g., in orange-fronted conures, (61), spectacled parrotlets, (34), or green-rumped parrotlets, (31)), and are presumed to do so in captivity (29). Consequently, the verbal interactions and exchanges of words and phrases with their human “flock” members are likely to represent the most socially relevant cues for parrots. Human words may be more difficult to imitate than other anthropogenic sounds (whistles or melodies) since they require finer tongue movement and articulation (12, 28). Thus, given the prolonged learning period related to longevity discussed above, it might not surprise that long-lived species are better at producing human words having prolonged association with human learning models.

Consistent with the higher probability of finding VPL in more gregarious species, we found that also the VPL total repertoire size was higher in more gregarious species. This finding aligns with the hypothesis that VPL of diverse repertoires may be adaptive when facing fluid social dynamics (62). It has been speculated that parrots’ highly social, fission–fusion foraging culture may be a result of their uniquely challenging diet of toxic, unripe seed, which may make social learning and information transfer about foraging sites (availability of riper and less toxic food, clay sources to neutralize toxins) indispensable (1, 13). Naive birds need to learn from individuals with knowledge of clay sources (geophagy to neutralize the toxins) and foraging sites (13). This social-learning-dependent system promotes strong social bonds within a group and encourages interaction with as many individuals as possible to gather information about finding food. This leads to flocks with overlapping areas where they search for food and frequently exchange individuals with different knowledge (1, 13). Interestingly, we found low intraspecific variation in VPL total repertoire size; in other words, species was a good predictor of VPL total repertoire size, i.e., of how many distinct mimicked vocalizations (different human words or anthropogenic sounds/melodies) a species exhibited. This consistency in VPL total repertoire size suggests a potential cap on memory capacity within species, implying that species exhibiting larger total repertoire in our dataset may inherently rely more on vocal mimicry in their natural environments. This could reflect an advanced cognitive ability to store and recall diverse vocalizations, which could be crucial for managing complex social interactions within their communities. However, it is important to note that while the repertoire size obtained in our survey showcases the parrots’ capacity for vocal learning, it does not necessarily translate to functional repertoire size in wild settings. The varied vocalizations likely represent adaptability to local conditions and interactions with humans and other species, rather than serving uniform functional roles. The extensive variety of mimicked sounds suggests that these parrots can adapt their VPL to suit unique environmental stimuli and social contexts, showcasing their vocal flexibility.

Finally, in accordance with our predictions, larger species were more likely to possess VPL. One potential explanation for our finding may be that large-sized parrots such as cockatoos and macaws but also amazons and grey parrots are typically housed as single individuals rather than in groups with conspecific, increasing the likelihood that they form strong bonds with their human caretakers and learn anthropogenic sounds vocalizations as part of their social interaction within their human “flock” (29). Furthermore, body mass shapes certain acoustic features of parrots’ vocalizations (e.g., peak frequency and duration) (37), potentially affecting the vocal plasticity of the differently sized species. In fact, we found that larger species had also larger VPL repertoires (including various anthropogenic sounds, melodies, and clearly distinguishable words). While small parrotlets and parakeets showed limited VPL repertoires ranging between 1 and 4 distinct vocalizations, larger-sized *Amazona* parrots, African grey parrots, cockatoos, and macaws tended to have larger VPL repertoires of up to 23 distinct learned vocalizations. This finding aligns with the neurological correlates of VPL described to date (10). For example, budgerigar, peach-faced lovebirds, and cockatiels, generally considered more limited in their VPL abilities, have larger and conspicuous core nuclei relative to the shell nuclei. In contrast, those thought to have considerably more complex communication abilities, such as the blue and gold macaws, peach-fronted conures, African grey parrots, and *Amazon* sp., have noticeably smaller cores and correspondingly larger shell regions, a parrot unique brain structure related to their advanced VPL capacities (10). Bigger species may exhibit greater vocal plasticity, but it is also likely that a large body size makes it easier to imitate the (low) pitch of human vocalizations compared to smaller species. Given that we used the imitation of human words as proxy for measuring differences in “VPL anthropophonic repertoire size”, we might have underrated the imitation complexity of small species, yet for determining the existence of VPL, any kind of allospecific sounds were used (including high frequency sounds, such as whistles, phone ring sounds etc.) thus providing a reliable indicator of whether a species is a vocal production learner or not.

In contrast to our predictions, relative brain size did not clearly affect VPL presence (neither the VPL total repertoire size nor VPL anthropophonic repertoire size). It is therefore possible that the cognitive demands of vocal learning/imitation (2, 41) have been overestimated or that relative brain size is a very crude proxy for the cognitive capacity required for VPL, as has been pointed out by several authors (43, 63). The density of forebrain neurons that may accommodate additional song and speech brain circuits has been proposed as a promising candidate for an alternative proxy (42).

### Conclusions and future directions

Vocal production learning (VPL) was likely an ancestral trait in parrots, present even in the most basal species. In due consideration of methodological challenges, our study reveals that VPL in parrots exhibits a complex evolutionary pattern of repeated gains and losses, emphasizing its adaptive value and the significant costs associated with its maintenance. Our findings also suggest significant relationships between VPL and key factors like social complexity, longevity, and body size. These elements shape the sophistication of vocal mimicry abilities in parrots, with larger, longer-lived species developing more complex vocalization strategies due to extended learning periods and social interactions. This pattern reflects a dynamic interplay between evolutionary pressures and the ecological niches occupied by these species, offering a unique lens into how cognitive abilities such as learning and memory interact with physical traits like brain and vocal tract morphology over evolutionary timescales. Exploring these changes can illuminate broader mechanisms of adaptability and resilience across different species.

For instance, investigating the genetic and neurological bases of VPL could elucidate how certain brain structures associated with vocal learning are preserved or altered amidst environmental pressures and genetic drift. Similarly, examining changes in vocal tract morphology and their correlation with vocal learning abilities could provide insights into the physical constraints and capabilities of parrots as they evolved. Additionally, parrots’ ability to reacquire VPL after losing it may help us understand the conditions under which complex traits can re-emerge in a lineage, offering broader implications for evolutionary theory. This makes the parrot a compelling model system for studying not just vocal production learning but also the evolutionary processes that govern the development and loss of complex behaviors more generally. Ultimately, these insights could spur new research into the evolutionary origins of human language, providing a valuable comparative framework to explore how complex communication systems have developed across different evolutionary pathways. This broadened perspective could significantly enhance our understanding of the mechanisms that underpin not only animal communication but also human language.

## Material and Methods

### Data collection

Data was compiled from the online video sharing and social media platform YouTube. We searched for scientific and common names of each parrot species, combined with words such as “talk”, “imitate”, or “sing” in two languages, English and Spanish (see Supplementary Information A, Table SA1 for further details). Species identified were visually confirmed, and scientific and common names were standardized using the IOC Bird List version 10.2 (64). Data were collected for all 398 extant parrot species (for a complete data set, see Github repository). We examined the first 50 hits for in- and exclusion criteria for each search. For a detailed description of the YouTube settings and in- and exclusion criteria, see Supplementary Information A.

In order to account for the likelihood of species being kept in human care (and thus be recorded on video when showing conspicuous behavior), we recorded for each species whether it was listed as being traded internationally. From the CITES homepage (https://trade.cites.org/) we obtained which of the 398 parrot species were categorized as “traded live” and listed them as “reported in trade”. We also compiled data on each species’ status in “captivity” (including on a regional level) from supplementary literature (65)(Reinschmidt 2006) and verified/complemented the information whenever possible by additionally consulting the Cornell Lab of Ornithology/Birds of the World homepage (https://birdsoftheworld.org/. Accordingly, we categorized the status in captivity as “common”, “rare”, and “not found” (full data set, GitHub repository). It was important to consider both variables because a species could be “common” in captivity, e.g., in a particular region where the parrots occur, but not be traded in the rest of the world. Vice versa, it could also be the case that, e.g., a critically endangered species might occur in international trade (because being part of breeding programs for conservation purposes) but remain restricted to few specialized breeders or facilities rather than being common in human care.

### Video analysis

After evaluation for in- and exclusion criteria, we (MS, JC, EH) and two extra coders blind to our research question analyzed videos that reached our inclusion criteria. For species with less than ten videos found, all videos were analyzed. In case ten or more videos were found, we analyzed at least ten videos per species (note that for 12 species *Agapornis fischeri, A. roseicollis, Amazona aestiva, A. albifrons, A. amazonica, A. auropalliata, A. autumnalis, A. farinosa, A. finschi, A. ochrocephala, A. viridigenalis, Anodorhynchus hyacinthinus* all videos found were analyzed ranging from 11 to 62 videos). The order of the search terms we used was always the same for each species. For all species for which videos with evidence for VPL (as described above) were found (163 species), the following variables were collected: 1) total number of mimicked vocalizations produced in the video (including repetitions), 2) template presence (yes/no), 3) quality of mimicked vocalizations (low/medium/high), 4) time bird is visible (seconds), 5) type of mimicked vocalization (i.e., a) anthropogenic non-lingual sounds, b) melodies, c) clearly distinguishable words or d) human-speech like babbling), 6) number of distinct mimicked vocalizations (i.e., how many different sounds/word were recognizable), 7) number of mimicked vocalizations for each type of vocalizations, 8) context in which the VPL occurred (i.e., whether the bird was mimicking while a) being alone, or b) in interaction with a) one or more human(s) or b) conspecific(s) or c) a combination of both), 9) language wherever possible to recognize (for all variables and relevant descriptions see Supplementary Information A, Table SA2). If multiple individuals of the same species appeared in one video and expressed mimicked vocalizations, the first one who started vocalizing was defined as the subject and was analyzed.

Note that the variables “total number of mimicked vocalizations (1)” and “language (9)” were not used in the analyses. The variable “number of distinct mimicked vocalizations (6)” was used to calculate the “VPL total repertoire size”. We took the number of distinct human words only to quantify the “VPL antropophonetic repertoire size”. Finally, the variable “quality of mimicked vocalizations” was used as “VLP quality” measure. The remaining variables (template, time, and context) were included in the analyses to control for confounding factors (for further description see Supplementary Information C, Figure SC1 Directed Acyclic Graph).

### Phylogeny

We used the phylogenetic supertree for all extant parrots presented recently by (66) to map the occurrence of VPL and tested whether the occurrence of VPL has a random distribution phylogenetically or is conserved in the phylogeny. The discrete character of VPL was mapped onto the phylogeny as a basic binary value (present or absent).

### Socioecological variables

Given the scarcity of information on social parameters for many parrot species we used data on gregariousness as a proxy for social complexity. The data on gregariousness were taken from (67). Gregariousness was scored as a categorical variable (1 = ‘yes’ or 2 = ‘no’) according to information from the ‘breeding’ section of the HBW Alive (68). A species was classified as gregarious if the description suggested that the breeding pairs nest close together or if the species was described as colonial. For details on data collection on longevity, body size, and relative brain size, see Smeele et al. (43).

### Data analysis

Our first goal was to map our findings/the occurrence of VPL in parrot species onto the phylogenetic tree of parrots (in order to reconstruct its evolutionary pathway within the order). Our second objective was to examine which variables best explained the presence of VPL across the parrot order. We ran ancestral state reconstruction with the function *contMap* from the package *phytools* (69). We also ran a Bayesian model to test the strength of the phylogenetic model. We modeled the probability of a species being able of VPL and included a variance-covariance matrix with the covariance as a function of the phylogenetic distance using the L2-norm.

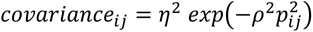

Where η and ⍴ are the parameters for the L2-norm and P is the normalized phylogenetic distance between species i and j (only for cases where i≠j). We used the L2-norm, since it does not assume Brownian motion, and allows for a non-linear decline in covariance with distance. We used Stan to define the models (70) and the package *cmdstanr* for model fitting (71) with four chains, 1000 burn-in and 1000 sample iterations, and *adapt_delta = 0.95*. We used R-hat to monitor model convergence and report any deviation outside 0.99-1.01 in the results section.

We were interested in testing the effect of longevity, relative brain size, sociality and body size on the probability that a species was capable of VPL. To ensure that correlations between these variables were not spurious, we included several covariates to the models. We used a Directed Acyclic Graph (DAG) to visualize the assumed causal paths (arrows) (72), see Supplementary Information C, Figure SC1). We then created a model for each variable of interest and used the back- door criterion (73) to determine which other variables to include as de-confounders. The back-door criterion ensures that only arrows pointing from the predictor variable towards the response variable (be it direct or indirect) are included in the parameter estimate. De-confounders are those variables that can create spurious correlations, when not controlled for (they point to the back-door of the predictor variable). All models converged well. The Rhat value of the etha^2 parameter and some of the intermediate species values on the log-odds scale (q) in the phylogenetic model were ∼1.01. The same was true for the intercept in the relative brain size model within the longevity model for the subset that only included pet species and rho^2 (∼1.02), etha^2 (∼1.01) and some q parameters in the phylogenetic model for this subset. All other Rhat values were within 0.99-1.01. We judged that these deviations were within the expected range of Rhat values and should not bias the results.

### Longevity

We hypothesized that longevity might affect the presence of VPL since longer-lived species may need to learn more new vocalizations throughout their lives due to their longer lives and potential exposure to more changes; also, long life spans may entail more opportunities for vocal learning and practicing. We included relative brain size, sociality, and body mass as covariates. We also included a varying effect for the genus to control for potential shared evolutionary history. Since our model contained missing values and we were interested in the effect of relative rather than absolute brain size, we ran a structural equation model similar to that suggested by Smeele et al. (43).

We modeled the presence of VPL with a binomial model (logit link) with the following formula:

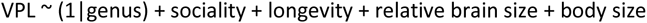

For a detailed model with all priors see Supplementary Information D.

#### Relative brain size

We hypothesized that relative brain size could affect the presence of VPL since VPL has been suggested to be cognitively demanding (10). We included sociality as a covariate, and the model structure was the same as that of longevity.

#### Gregariousness

We hypothesized that gregariousness may affect the presence of VPL since VPL can have social functions and therefore gregarious species might be relying on using learnt vocalizations more than less gregarious species and may also have more opportunities or greater need to learn new ones socially. We did not include any covariates, and the model structure was the same as that of longevity.

#### Body size

We hypothesized that body size might affect the presence of VPL. Larger species have a larger syrinx, which might make mimicry of the human voice and speech more likely. Furthermore, we expected larger species to be more often housed as single individuals, increasing the likelihood that they mimic the vocalizations of their human caretakers (29). We did not include any covariates. The model structure was the same as that of longevity.

#### Interspecific variation in VPL total repertoire size and VPL antropophonic repertoire size

The third goal of our study was to understand what factors drive interspecific variation in VPL, assessing both total repertoire size and anthropophonic repertoire size. Therefore, we scored the overall number of distinct mimicked vocalizations (i.e., different learnt vocalizations) and the number of distinct human words only produced by different species. We did not include the duration of the recording (i.e., the time the bird was visible) in the model, because this variable did not create a backdoor path between the variables of interest (longevity, sociality, body size, and relative brain size) and the response variable (number of distinct mimicked vocalizations). In our DAG, the duration only affected whether or not a distinct mimicked vocalization was detected, but it did not affect the other variables and also was not affected by any variables. Therefore, the parameter estimates should not be biased by duration. We modeled the VPL total repertoire size with a Poisson model (log link) with the same formula as the VPL presence model. To analyze the interspecific variation in VPL total repertoire size, we had multiple observations, i.e., from different individuals per species. Therefore, we expanded the models to include the varying effect of species and genus. For a detailed model with all priors as well as the detailed hypotheses see Supplementary Information D.

### Ethical statement

Our methodology involved the use of publicly available online videos; no private or restricted content was accessed. We maintained the anonymity of all individuals and entities involved. No personal data was collected, and any potentially identifying information present in the videos was omitted from our analysis and reports to protect the privacy of individuals. We did not attempt to contact video uploaders or subjects, and we refrained from using data for purposes other than those explicitly stated in our research objectives. Consent for the use of these videos in a public domain was presumed based on their publicly accessible status and the nature of social media sharing. However, we acknowledge that the interpretations and analyses presented in this study are solely those of the research team and do not reflect the views of the video creators or subjects.

### Data accessibility

All data and code used in this study and all code to reproduce the analyses in this study are publicly available and maintained on GitHub: https://github.com/simeonqs/Insights_into_the_phylogenetic_distribution_and_evolution_of_vocal_production_learning_in_parrots. All data and code will also be permanently stored on Zenodo upon acceptance and can be accessed for review under: https://zenodo.org/records/14051008?preview=1&token=eyJhbGciOiJIUzUxMiJ9.eyJpZCI6Ijg0ZGI2MDE0LTY1NDktNDIzMy04NjJjLTQzMTFiYjcyNDVmOCIsImRhdGEiOnt9LCJyYW5kb20iOiI2ZTI2NTUwZmRkNzM2YTM0MWUwMmNhY2U2ZWZjMDY2OCJ9.NtmZyNyGQjEk-5GW7uTKRkuqYrxWHH-JNXWhuOaiN8V7iEz8iYro4I0OvPIXWWoPw0IKOArqSHcNI1l7gzXubw

## Acknowledgements

We thank the Loro Parque Fundación for their support and advice. We also thank Garance Barbier and Sayuri Diaz for assisting with data collection and scoring. EH received funding from DAAD, Germany in the GSSP program.

