## Supplementary material for "Evolution of vocal production learning in parrots": Supplemetary Information A-D

### Supplementary information A

*YouTube settings*

'Type' was set to 'Video'; videos were sorted for relevance. Browser history was cleared between species, and YouTube search and watch history was cleared between searches to minimize the possibility that the YT recommendation algorithm affects the shown results based on the previous search history.

*In- and exclusion criteria*

For a video to be included, the title had to include the word 'talk(ing)', 'imitat(ing)' or 'sing(ing)', 'laugh(ing)' or a conjugation or synonym in the languages English and Spanish (see Supplementary Information Table SX4 for exact keywords and conjugations used). This was done to include only relevant videos for analysis, namely videos with a high chance of VPL occurrence.

The following exclusion criteria were applied to keep the data scoring feasible: (1) videos over 20 minutes long, (2) videos, in which it was not possible to determine the species within the first 20 seconds of it being in view due to either camera quality, lighting, or distance from the camera, (3) videos, in which the first parrots shown were four or more individuals simultaneously in the frame for at least 20 seconds and (4) videos that did not feature any parrot within the first 15 seconds. In addition, to ensure that the sounds hearable in the video are actually produced by a parrot we excluded: (5) videos, in which no parrot was clearly in frame and vocalizing within the first 50 seconds it had first appeared, and (6) videos in which further species featured after a first species was shown for at least 10 seconds. Further, any type of edited videos, such as (7) videos edited for informative purposes, (8) videos featuring still images (from the start of the video for at least 10 seconds), (9) documentary-style videos (voiceover for at least 10 seconds), (10) videos with added music (at least 15 seconds from the start of the video), (11) tutorial-style videos on teaching a parrot how to talk, and (12) compilations of several videos, were excluded to ensure the behaviors being observed were not edited or manipulated.

When multiple videos from the same YouTube user were found for a given species, only one was analyzed, namely, the first one found.

*Species evaluation*

For each video, we evaluated if the indicated species was indeed shown by making a visual assessment using the identification guide ebird.org (Cornell Lab of Ornithology). When the species featured was not clearly identifiable, the video was excluded.

**Table SA1.** Search terms used in each YouTube search and their conjugations (in brackets), as well as the synonyms of search terms that were used as video inclusion criteria (including their conjugation, in brackets).

| **English search terms** | **Synonyms** | **Spanish search terms** | **Synonyms** |
|---|---|---|---|
| Talks (talk*) | Speaking (speak*), vocalizing (vocalis*), saying (say*), conversing (convers*), discussing (discuss*), chatting (chat*) | Habla (habl*) | Conversar (convers*), decir (dig*, dice*, dicho, deci*, dir*, dij*, di) |
| Imitates (imitat*) | Mimicking (mimick*), copying (copy*), impersonating (impersonat*) | Imita (habl*) | Simular (simul*), copiar (copi*) |
| Sings (sing*) | Whistling (whistl*) | Canta (cant*) | Silbar (silb*) |
| Laughs (laugh*) | NA | Ríe (rei*) | NA |

**Table SA2.** Variables recorded for every video, with corresponding levels and descriptions when needed.

| Variable | Levels | Description |
|---|---|---|
| Vocal learning presence | Yes, no | Production of clearly allospecific or environmental sounds (such as human words, ringing bells, melodies). |
| Template presence | Yes, no | Within 60 seconds before the recorded imitation, a template sound was heard. |
| Quality of mimicked vocalizations | Low, moderate, high | **Low:** No recognizable words, only melodic imitation<br>**Moderate:** Barely distinguishable word(s)<br>**High:** Very clearly distinguishable word(s) |
| Time bird is visible | seconds | The duration that the bird is in the frame during the video |
| Type of sound | Human word(s), human sound, human babbling, singing, allospecific nonhuman sound, environmental sound | **Human sound:** For example, laughter or another human sound that does not involve words.<br>**Human babbling:** Indistinguishable as words but distinguishable as human-like vocalizations.<br>**Singing:** Imitation of a melody or a human whistle.<br>**Environmental sound:** For example, a phone ringtone or the sound of water. |
| Number of distinctive imitations | - | Any clear word, sentence, melody or other distinctive sound without any interruption longer than 3sec was counted as a distinct imitation. An uninterrupted sequence of human babbling in a video was counted as one distinct imitation. |
| Number of imitations for types of sounds | - | For each type of sound, the number of distinct imitations was noted. For example, when a bird said two distinctive words, two imitations were noted for 'human words'. |
| Context | Alone, interaction with humans, with conspecifics, with nonhuman allospecifics | **Interaction with humans:** The bird is either in close proximity with a human, or the human is visibly or audibly engaging with the bird.<br>**With conspecifics:** The bird is in close proximity to other parrots.<br>**With nonhuman allospecifics:** Another animal, e.g., a dog, is visible in the video |
| Language | - | The language of a template human word/song, if applicable. |

### Supplementary information B

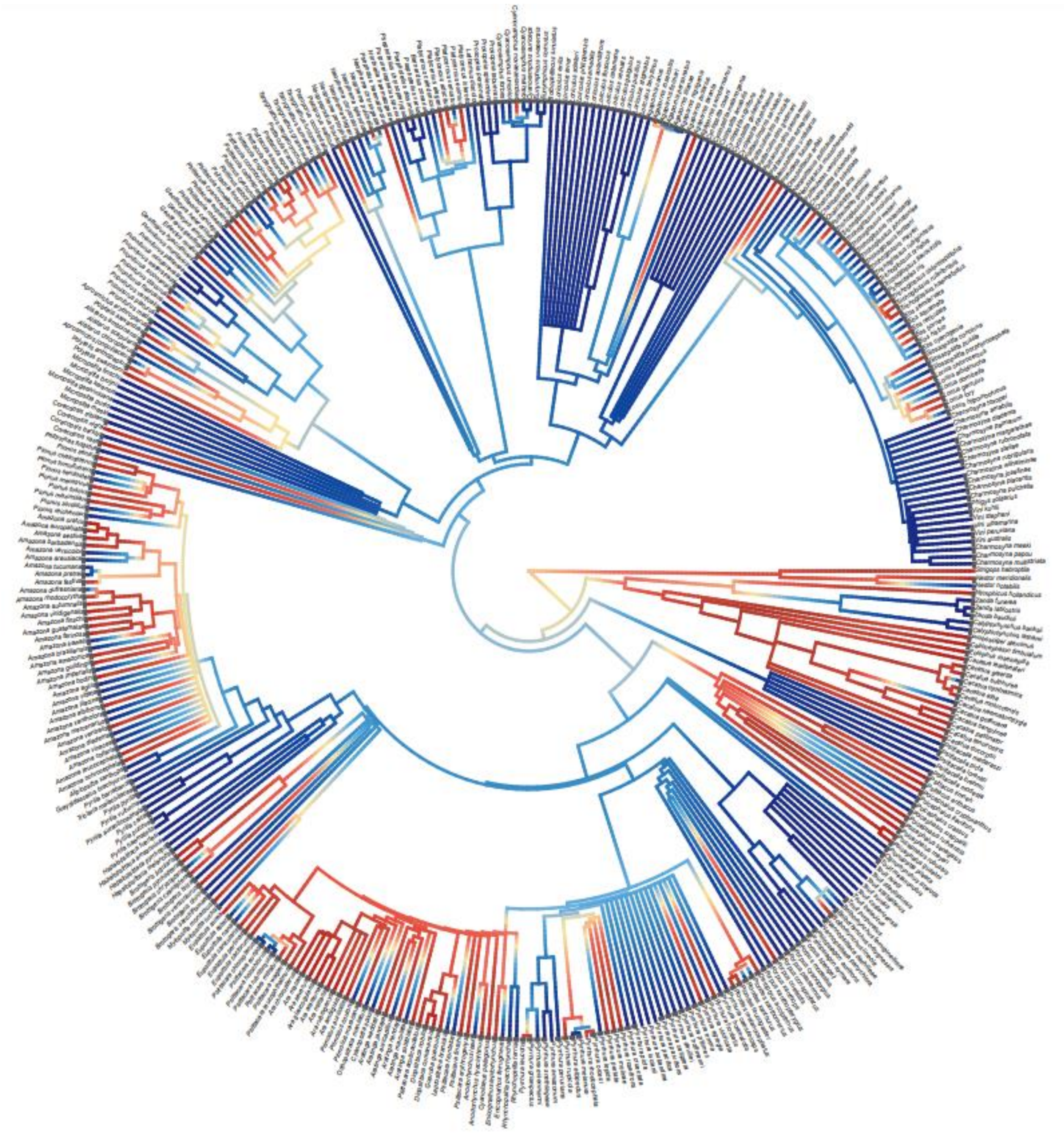

**Figure SB1:** Phylogenetic tree of all parrot species on the species level indicating presence or absence of VPL in a species or lack of data on their VPL ability. Colors represent a conservative ancestral state reconstruction of VPL ranging from blue = no VPL detected to red = VPL detected. The grey squares indicate the species found in human care. Missing grey squares indicate the 44 species that are not found in human care. Note that for 43 out of these 44 species, VPL ability remains unknown (=missing data points).

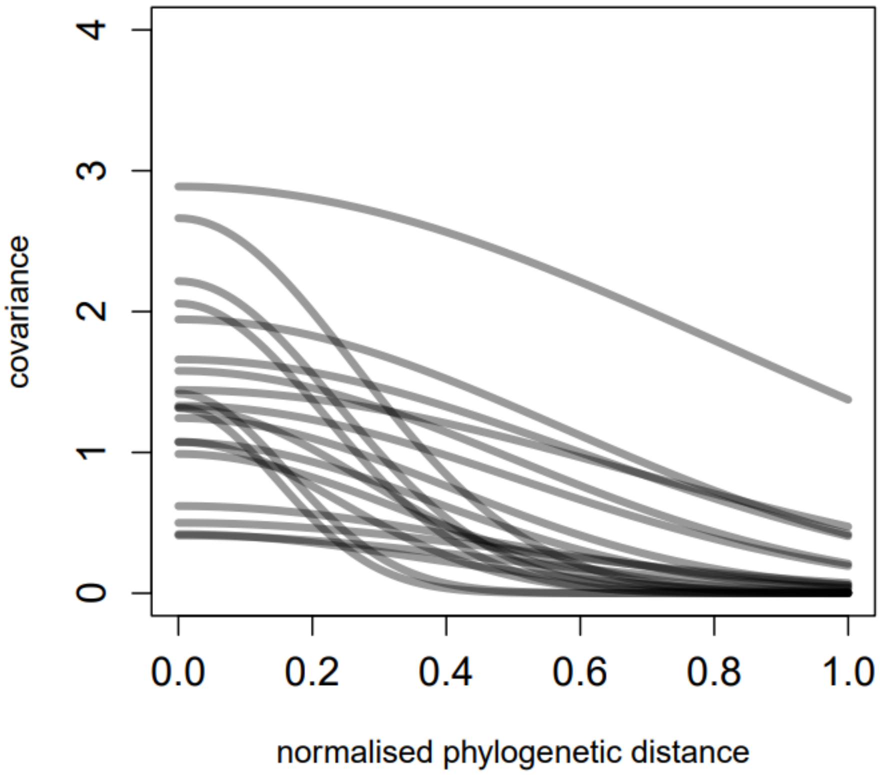


**Figure SB2:** Strength of the phylogenetic model based on a Bayesian model for the probability of a species being a VPL with the covariance as a function of the phylogenetic distance using the L2-norm: $covariance_{ij} = \eta^2 \; exp\left(-\rho^2 p_{ij}^2\right)$

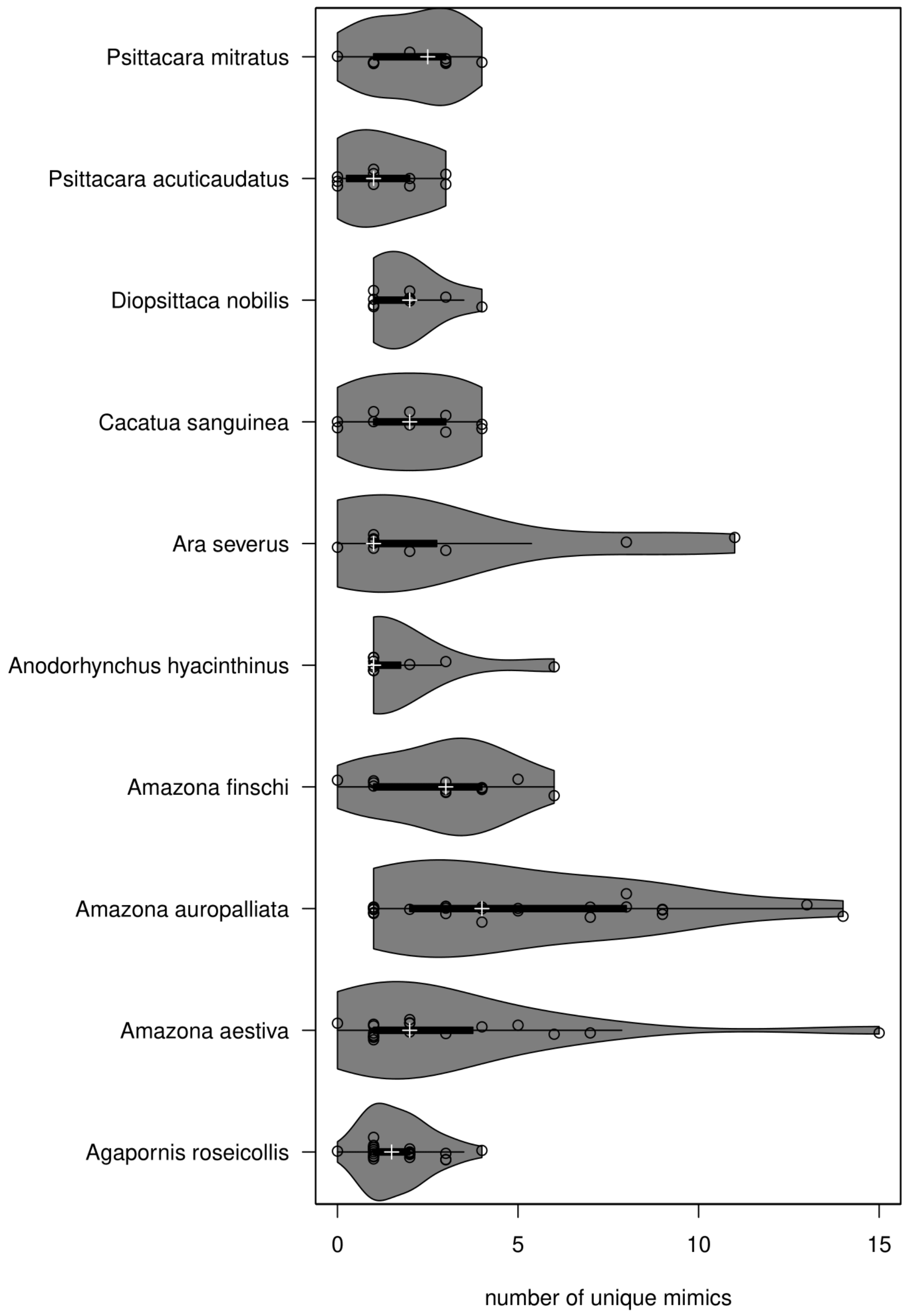


**Figure SB3:** Within-species variation for 10 randomly chosen species (with at least 10 videos of different individuals per species available). Circles are raw data, thin and thick black lines are boxplots that exclude outliers, grey shaded areas are violin plots and white crosses are means.

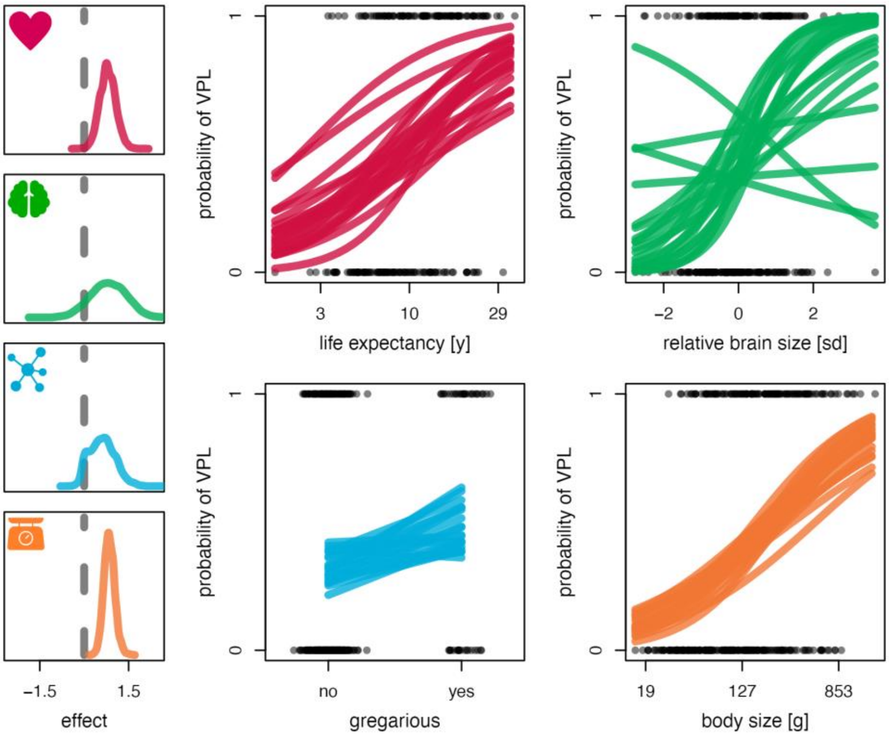


**Figure SB4:** Variables influencing a species' probability of exhibiting VPL (including only species reported in human care). Lefthand side: posterior densities of the effect of longevity (red), relative brain size (green), gregariousness (blue), and body size (orange). For gregariousness, the contrast between a non-gregarious and a gregarious species is shown. For all other variables, the slope is shown. Righthand side: scatterplots of the raw data (grey) and 20 posterior predictions (colored lines) per variable.

**Table SB1.** Effect of longevity, sociality, relative brain size, and body size on VPL quality

| effect | mean | Posterior intervals (PI) |
|---|---|---|
| total effect longevity (beta) | 0.41 | 0.07-0.75 |
| total effect sociality (contrast) | 0.16 | -0.16-0.61 |
| total effect relative brain size (beta) | 0.08 | -0.99-0.83 |
| total effect body size (beta) | 0.88 | 0.65-1.1 |
| total effect template (contrast) | 0.01 | -0.25-0.26 |
| sigma effect context | 0.27 | 0.02-0.73 |

### Supplementary information C

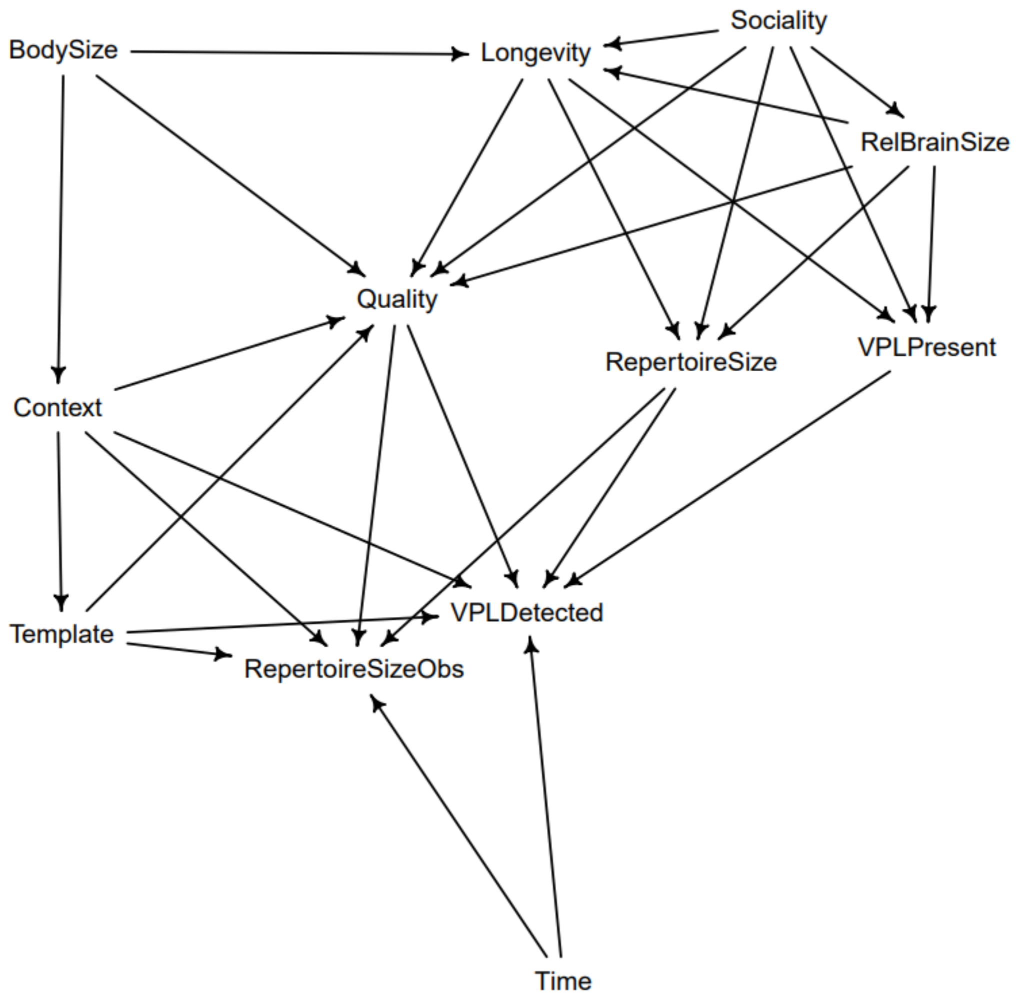


**Figure SC1:** Directed Acyclic Graph (DAG) to determine which covariates to include in each model. All assumed causal relationships between variables are represented with arrows. Each arrow represents the direction of causality. Following variables were taken into consideration: *VPLPresent*=occurrence of VPL (yes/no), *VPLdetected* = whether or not VPL is observed in a YouTube video, *RepertoireSize* = number of distinct mimicked vocalizations an individual produced, *RepertoireSizeObs* = the observed number of distinct mimicked vocalizations in the video (assuming that an individual is unlikely to produce everything it knows; used to check if it was necessary to 'control' for duration of the video). Time= the duration that the subject is visible in the video (video duration ranged from 4s to 897s; mean±SD: 72±111s); Template=presence of a template for the mimicked vocalization (produced either by human or other anthropogenic source). Context = context in which the VPL was detected (subject being alone / interacting with human / and/or conspecifics; Quality = assessment of the recognisability of the mimicked vocalizations (low/medium/high); RelBrainSize= relative brain size (for data extraction see Smeele et al. (2022); Body size=body mass based on data obtained from ZIMS (for data extraction see Smeele et al. (2022); Longevity= average life expectancy based on obtained data on birth and death dates from Species360's ZIMS (Smeele et al. 2022); Sociality = gregariousness scored as a categorical variable (1 = 'yes' or 2 = 'no') according to information from the 'breeding' section of the HBW Alive (Del Hoyo et al. 2017; for data extraction see Carballo et al. (2020).

### Supplementary information D

Model for the probability of a species being able of VPL. The covariance between any two species was a function of the phylogenetic distance:

$$M \sim binomial(1,\ p)\ M = mimicry\ or\ not\ logit(p)\ \sim Mvnormal(\mu,\ S)\ p$$
$$= probability\ of\ mimicry\ \ \mu\ =\ average\ log - odds\ mimicry\ \ S$$
$$=\ variance\ covariance\ matrix\ S_{ij}$$
$$= \eta^2\ exp\left(-\rho^2 P_{ij}^2\right)\ +\ \delta_{Pij}\ \sigma_P^2\ formula\ for\ L2\ norm\ \eta^2\ \sim exponential(2)\ P$$
$$=\ normalized\ phylogenetic\ distance\ \rho^2\ \sim exponential(0.1)\ \eta^2, \rho^2$$
$$= parameters\ for\ L2\ norm$$

The main model for the presence of VPL was as follows:

$$model\!:_{\_}\ M\ \sim binomial(1,\ p)\ M = mimicry\ or\ not\ logit(p)\ = \bar{\alpha}\ + \alpha_G\ + \alpha_S\ +\ p$$
$$= probability\ of\ mimicry\qquad \beta_L * L\ + \beta_{RB} * RB\ + \beta_B * B\qquad priors\!:_{\_}\ \bar{\alpha}$$
$$\sim normal(0,\ 1)\ average\ log - odds\ mimicry\ \alpha_G$$
$$\sim normal(0,\ \sigma_G)\ off - set\ for\ genus\ \sigma_G$$
$$\sim exponential(2)\ between\ genera\ standard\ deviation\ \alpha_S$$
$$\sim normal(0,\ \sigma_S)\ off - set\ for\ sociality\ \sigma_S$$
$$\sim exponential(2)\ between\ sociality\ standard\ deviation\ \beta_L$$
$$\sim normal(0,\ 1)\ slope\ for\ longevity\ \beta_{RB}$$
$$\sim normal(0,\ 1)\ slope\ for\ relative\ brain\ size\ \beta_B$$
$$\sim normal(0,\ 1)\ slope\ for\ body\ size$$

Missing values for longevity were imputed with the following sub-model:

$$model\!:_{\_}\ L\ \sim normal\left(\mu_L,\ \kappa_L\right)\ L = standardized\ life\ expectancy\ \mu_L$$
$$= \bar{\theta}\ + \theta_G\ + \xi_{RB} * RB\ + \xi_B * B\ \mu_L = average\ life\ expectancy\qquad priors\!:_{\_}\ \kappa_L$$
$$\sim exponential(2)\ standard\ deviation\ life\ expectancy\ \bar{\theta}$$
$$\sim normal(0,\ 1)\ average\ life\ expectancy\ \theta_G$$
$$\sim normal(0,\ \kappa_G)\ off - set\ for\ genus\ \kappa_G$$
$$\sim exponential(2)\ between\ genera\ standard\ deviation\ \xi_{RB}$$
$$\sim normal(0,\ 0.5)\ slope\ for\ relative\ brain\ size\ \xi_B$$
$$\sim normal(0,\ 0.5)\ slope\ for\ body\ size$$

Relative brain size ($RB$) was computed, and missing values were imputed with the following sub-model:

$$model\text{:}\_\ Br \sim normal(\mu_B,\ \phi_B)\ Be = standardized\ brain\ size\ \mu_B\ = \bar{\omega} + \omega_G\ + \gamma_B * B\ \mu_B$$
$$= average\ brain\ size\ PB\ = \bar{\omega} + \gamma_B * B\ PB = predicted\ brain\ size\ RB$$
$$= B - PB\ RB = relative\ brain\ size\quad priors\text{:}\_\ \phi_B$$
$$\sim exponential(2)\ standard\ deviation\ brain\ size\ \bar{\omega}$$
$$\sim normal(0,\ 1)\ average\ brain\ size\ \omega_G$$
$$\sim normal(0,\ \phi_G)\ off - set\ for\ genus\ \phi_G$$
$$\sim exponential(2)\ between\ genera\ standard\ deviation\ \gamma_B$$
$$\sim normal(0,\ 0.5)\ slope\ for\ body\ size$$

*Model for the total repertoire size of VPL*

*Longevity*

Again, we hypothesized that longevity could affect VPL total repertoire size (in terms of overall number of distinct mimicked vocalizations) since longer-lived species may employ learnt vocalizations more often and have more time to learn new vocalizations. The main model was as follows:

$$model\text{:}\_\ NM \sim poison(exp(\lambda))\ NM = number\ of\ mimics\ \lambda\ = \bar{\alpha} + \alpha_G\ + \alpha_{Sp} + \alpha_S\ +\ \lambda$$
$$= average\ number\ mimics\quad \beta_L * L\ + \beta_{RB} * RB\ + \beta_B * B\quad priors\text{:}\_\ \bar{\alpha}$$
$$\sim normal(2,\ 2)\ global\ average\ number\ mimics\ \alpha_G$$
$$\sim normal(0,\ \sigma_G)\ off - set\ for\ genus\ \sigma_G$$
$$\sim exponential(2)\ between\ genera\ standard\ deviation\ \alpha_{So}$$
$$\sim normal(0,\ \sigma_{So})\ off - set\ for\ species\ \sigma_{So}$$
$$\sim exponential(2)\ between\ species\ standard\ deviation\ \alpha_S$$
$$\sim normal(0,\ \sigma_S)\ off - set\ for\ sociality\ \sigma_S$$
$$\sim exponential(2)\ between\ sociality\ standard\ deviation\ \beta_L$$
$$\sim normal(0,\ 1)\ slope\ for\ longevity\ \beta_{RB}$$
$$\sim normal(0,\ 1)\ slope\ for\ relative\ brain\ size\ \beta_B$$
$$\sim normal(0,\ 1)\ slope\ for\ body\ size$$

The sub-models to impute longevity and calculate relative brain size were the same as the model of VPL presence. It should be noted that longevity and relative brain size were imputed at the species level, while the main model contained multiple observations from different individuals for some species.

*Relative brain size*

Relative brain size might affect the VPL total repertoire size (regarding the number of distinct mimicked vocalizations) since larger brains would allow a species to learn and remember more distinct mimicked vocalizations. We included sociality as a covariate. The model structure was the same as that of longevity.

*Sociality*

We hypothesized that sociality might affect the VPL total repertoire size (i.e., the number of distinct mimicked vocalizations) since social species would benefit more and would have more opportunities to learn new vocalizations through social learning. We did not include any covariates. The model structure was the same as that of longevity.

*Body size*

We hypothesized that body size could affect the VPL total repertoire size (i.e., the number of distinct mimicked vocalizations) since larger species have a larger syrinx, making mimics of human voices/speech more likely. Furthermore, we expected larger species to be more often housed as single individuals, increasing the likelihood that they mimic the vocalizations of their caretakers. We did not include any covariates. The model structure was the same as that of longevity.

The same models were used to test for those effects also on the number of distinct mimicked human words only, i.e. VPL anthropophonic repertoire size.
